# Genome wide analysis in *Drosophila* reveals diet by gene interactions and uncovers diet-responsive genes

**DOI:** 10.1101/718304

**Authors:** Deanne Francis, Shila Ghazanfar, Essi Havula, James R. Krycer, Alistair Senior, Annabel Y. Minard, Thomas Geddes, Fiona Weiss, Jacqueline Stöckli, Jean Y.H. Yang, David E. James

**Author notes:** Corresponding author: David E. James, The University of Sydney, Camperdown, 2006, New South Wales, Australia. Author contributions: D.F. and D.E.J. conceived the study. D.F., J.R.K and D.E.J participated in experimental design. D.F., F.W., and A.M. performed the experiments and D.F., A.S., E.H., T.G. and S.G. analysed data for the experiments. J.S. and J.Y. provided intellectual input. D.F., J.S. and D.E.J. wrote the manuscript, and all the authors edited the manuscript. D.E.J. supervised the study.

## Abstract

Genetic and environmental factors play a major role in metabolic health. However, they do not act in isolation, as a change in an environmental factor such as diet may exert different effects based on an individual’s genotype. Here, we sought to understand how such gene-diet interactions influenced nutrient storage and utilisation, a major determinant of metabolic disease. We subjected the Drosophila Genetic Reference Panel (DGRP), comprising 200 genetically divergent inbred fly strains, to diets varying in sugar, fat and protein. We assessed starvation resistance, a holistic phenotype of nutrient storage and utilisation that can be robustly measured. Diet influenced the starvation resistance of each strain, but this effect varied markedly between strains. This demonstrates that genetics plays a major role in the response to diet. Furthermore, heritability analysis revealed that the greatest variability arose from diets either high in sugar or high in protein. To uncover the genetic underpinnings of this variation, we mapped 1,239 diet-responsive SNPs in 534 genes, 325 of which have human orthologues. Using whole-body knockdown, we confirmed that 30 candidate genes were required for glucose tolerance, storage and utilization. In particular, we characterised CG4607, a GLUT6/GLUT8 homolog, as a key protein involved in sugar tolerance. Overall, this provides strong evidence that genetics is a major contributor to how individuals respond to diets of varying nutrient composition. It is likely that a similar principle may be applied to metabolic disease in higher organisms thus supporting the case for nutrigenomics as an important health strategy.

## Introduction

Although it is widely appreciated that gene and environment interactions play a major role in determining various phenotypes in different animals, the application of this paradigm to health outcomes in humans remains an unmet need. For instance, a well-accepted culprit of metabolic disease is the widespread introduction of the high-fat, high-sugar ‘Western’ diet. Interestingly, indigenous populations only experienced a considerable burden of metabolic disease upon exposure to Western diets, after subsisting on different diets, whether rich in fats (Greenlandic Inuits) or carbohydrates (American Pima Indians), for thousands of years (Schulz & Chaudhari, 2015; Andersen & Hansen, 2018). However, studies in mice and humans have demonstrated that individuals display heterogeneous metabolic responses to the same diets (Zeevi *et al*, 2015; Parks *et al*, 2015). While genome-wide association studies (GWAS) of body mass index, insulin resistance and other metabolic traits have identified several causative loci, these loci do not account for the majority of phenotypic variation (Phillips, 2013). In fact, studies in mono- and di-zygotic twins revealed that a combination of genetic and environmental factors contributed to variance in body weight (Stunkard *et al.*; Dubois *et al.* 2012), with obesity being linked to metabolic disease risk. Overall, this emphasises the need to consider not only genetic predisposition or diet alone, but specifically gene-diet interactions (Heianza *et al.* 2017). Furthermore, one’s genotype can affect metabolic outcomes in response to a particular diet, and this ‘diet-responsiveness’ can vary markedly between individuals.

The notion that individuals display discrete sensitivity to certain macronutrients, with dietary interventions being based on one’s genotype, is the key premise to the field of personalised nutrition, or nutrigenomics (Sales *et al.* 2014) and is supported by recent preliminary clinical trials (Qi, 2014; Tan *et al*, 2019). However, we do not have a thorough understanding of how genes and diet interact to influence the risk of metabolic disease (Drabsch & Holzapfel, 2019). Here, we aimed to set up a model system for recapitulating gene-diet interactions. Studying this in humans remains a challenge because the environmental variables are difficult to control at a sufficient scale to facilitate genetic mapping. In contrast, the *Drosophila* fruit fly model system overcomes many of these logistical issues. Importantly, >70% of known human disease genes have fly orthologs (Reiter *et al.* 2001), and genetic tools such as the *Drosophila* genetics reference panel (DGRP) with 200 inbred and fully sequenced lines are available, thereby allowing identification of causal genetic variants (Mackay *et al.* 2012).

We capitalised on the high-throughput nature of the DGRP to study the effect of gene-diet interaction on nutrient storage and utilisation. A phenotype that encapsulates these outcomes is survival during starvation, which measures an organism’s ability to metabolise and store dietary nutrients during feeding and utilise them efficiently during fasting. For instance, adult flies fed a high carbohydrate diet showed greater resistance to starvation, as well as higher triacylglycerides (TAGs), suggesting a correlation between diets and starvation resistance via energy storage (Lee and Jang 2014) – however, does this apply to all individuals? Indeed, there was significant variation in starvation resistance between strains across the DGRP when fed a single diet (Mackay *et al.* 2012). Thus, we utilised the genetic diversity of the DGRP to identify SNPs that alter starvation resistance after exposure to diets varying in fat, sugar and protein content. Our study provided strong evidence establishing starvation resistance in *Drosophila* as a prime example of a phenotype governed by gene-diet interactions, motivating the notion that a similar principle may be applied to metabolic disease in higher organisms. Here, we combine the high-throughput nature of the *Drosophila* model with the genetic diversity of the DGRP to dissect diet-gene interactions on a population level. In this study, we aimed to identify “diet-responsive” genes and determine the mechanism by which genes in combination with diet control metabolic phenotypes. To do this, we used the DGRP to perform a GWAS to identify SNPs that contribute to variation in response to diets that differ in fat, sugar and protein contents. Our study uncovers a previously under-appreciated influence of diet on the heritability of starvation resistance. Finally, we provide a rich resource of diet specific genes for further study.

## Results

### Starvation Resistance of DGRP across 4 different diets

We first sought to identify novel diet-responsive genes that affect nutrient storage and utilisation in *Drosophila*. We used survival during starvation as a surrogate for an obesogenic phenotype to screen for dietary effects. This is a powerful and sensitive assay as starvation resistant flies are often replete with fat stores immediately prior to starvation and feeding flies a high sugar diet increases fat stores and prolongs starvation resistance (Djawdan *et al.* 1998; Hoffmann and Harshman 1999; Bjedov *et al.* 2010). The diets we selected, (normal food (NF), high carbohydrate diet (HCD), high fat diet (HFD) and high protein diet (HPD), Table S1), were based on previous studies that explored the effect of different sugar and protein concentrations on starvation resistance in a single strain (Skorupa *et al.* 2008; Lee and Jang 2014; Chandegra *et al.* 2017). The dietary composition is indicated in Table S1: the carbohydrate is sugar, the protein is yeast and the fat is coconut oil. We exposed 3 to 5-day old adult males from 178 DGRP strains to the 4 diets for 10 days and then measured starvation resistance by removing food and assessing survival (Fig. 1A, Fig. S7). The DGRP has previously been used to examine starvation resistance in flies fed NF (Mackay *et al.* 2012) and there was a strong correlation in starvation resistance across the 178 strains between the two studies (males, NF, Pearson’s R= 0.58, Fig. 1B), demonstrating the robustness of the starvation phenotype and the DGRP resource. Interestingly, we found that previously published food intake data (Garlapow *et al.* 2015) was negatively correlated with starvation resistance across genotypes (males, NF, Pearson’s R= −0.32, Fig. 1B) in both our study and previously published starvation data (Mackay *et al.* 2012). This is intriguing as it suggests that strains that ate the most were the least resistant to starvation. This could be due to differences in metabolic rate, an increase in hunger cues, nutrient storage capacity or differences in hormonal responses.

**Figure 1:**
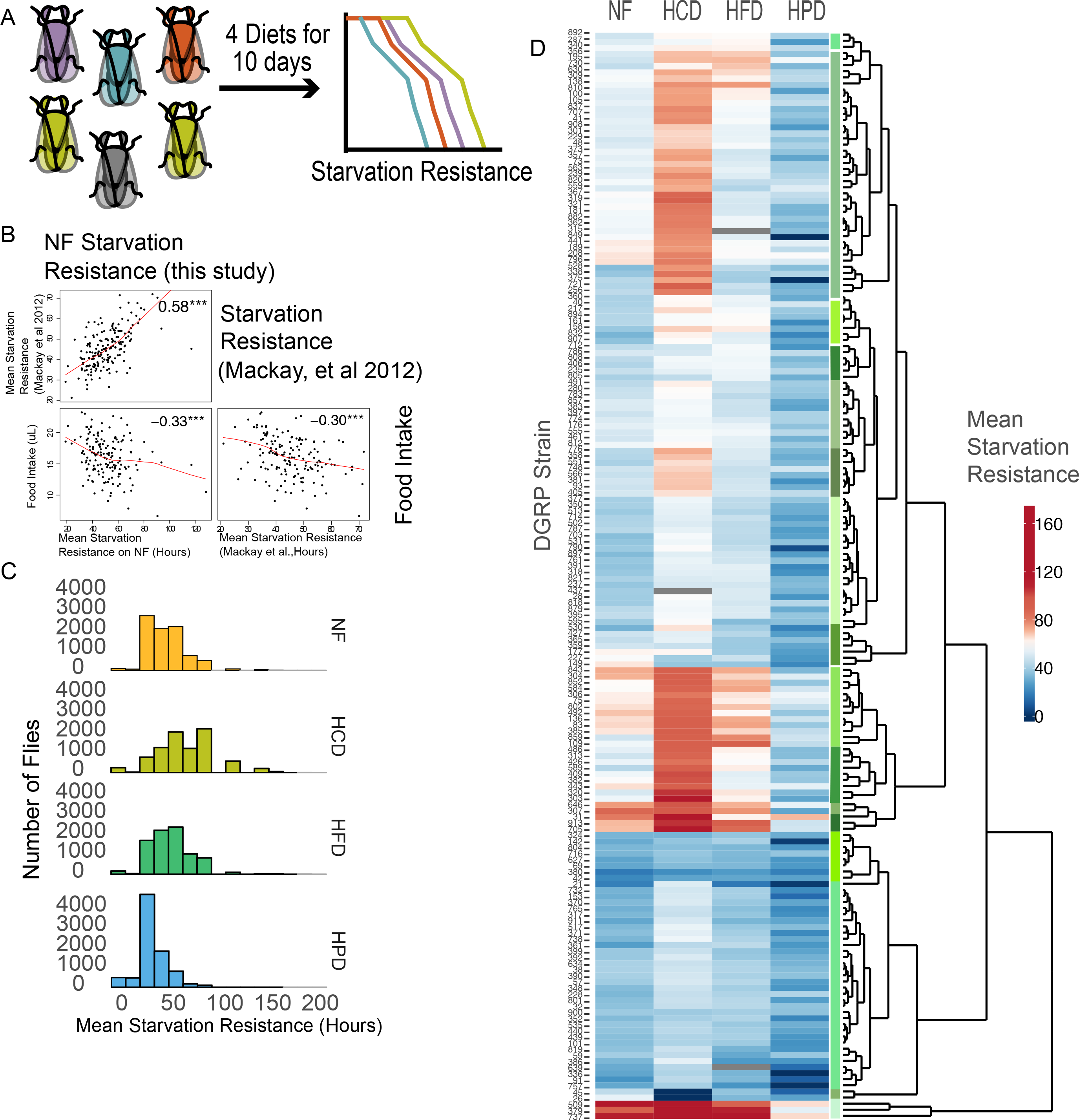
Inter-strain variation in response to diet. (A) DGRP screen schematic outlines the work flow. Each individually coloured fly represents an individual DGRP line and survival curves represent the measured starvation resistance after exposure of flies to each diet. (B) Pairs plot showing a positive correlation with starvation resistance data from Mackay et al, 2012 and negative correlation with food intake to starvation resistance data on NF from this study. (C) Histograms showing the distribution of the mean starvation resistance of each fly independent of strain on each diet. (D) Heatmap and hierarchical clustering of the mean starvation resistance of each DGRP strain.

Irrespective of strain, flies fed NF, HFD and HCD displayed the greatest variance in starvation resistance, while HPD fed flies had the least amount of variation across the strains (Fig 1C, 1D). We normalised the starvation resistance to NF for each strain and performed hierarchical clustering, indicating clusters of strains that were starvation sensitive to either HCD, HPD or HFD (Fig S1). Surprisingly these data showed a disparity between the responses to different diets, likely driven by genetic background, indicating that there is no single optimal diet where all strains display similar starvation resistance responses (Fig 1D, Fig S1).

We quantified the differential contributions of diet and gene by diet interactions to the variation in starvation resistance among DGRP strains. We determined the broad sense heritability (H_2_) of the starvation response using linear-mixed models (LMMs). We found that across the whole data set, after accounting for differences among diets, the H2 of starvation resistance was 20%. Dietary effects accounted for 8% of the variance in starvation-resistance, leaving 72% residual (unexplained) variance. A model including a term to estimate genetic variance in dietary effects (random-slopes LMM) had a significantly better fit than a model without (likelihood ratio test, *L* = 11611, d.f. = 14, p<0.001) indicating the presence of gene by environment interactions. Within-diet, the H2 of starvation resistance on NF, HCD, HFD and HPD was 19%, 50%, 20% and 65%, respectively (Table 1 and S2). Thus, across populations certain diets expose genetic diversity more strongly than others and we estimate that genetics plays a particularly strong role in determining starvation resistance in flies fed either HPD or HCD.

**Table 1:** Analysis of the mean starvation resistance data on all diets.

| **Diet** | **Mean** | **SEM** | **Variance**<br>**(Genetic)** | **Broad Sense**<br>**Heritability** |
| --- | --- | --- | --- | --- |
| **NF** | 51.46 | 1.15 | 0.086 | 19% |
| **HCD** | 68.88 | 1.64 | 0.364 | 50% |
| **HFD** | 55.10 | 1.09 | 0.089 | 20% |
| **HPD** | 36.77 | 0.97 | 0.671 | 65% |

A summary table of the mean starvation data, SEM, genetic and environmental variance from the log transformed starvation data across all the DGRP lines on each diet.
Estimates of genetic variance and heritability are derived from linear mixed models of log survival (see supplementary table S2).

### Mapping ‘diet-responsive genes’

We aimed to identify gene-diet interactions by uncovering SNPs associated with diet-responsive starvation resistance (Fig.2A). We first identified diet-associated SNPs across all diets by performing multivariate ANOVA testing with a significance threshold of unadjusted P-value < 1×10^−4^. This is higher than the conventional threshold used (P value 1× 10^−5^, (Mackay *et al.* 2012)) to increase the number of true positive SNPs as further filtering was applied. Such filtering reduced the number of SNPs to exclusively contain SNPs that exhibited a significant and large difference in starvation survival compared to NF in at least one diet (unadjusted p-value < 0.01, absolute log fold change > 0.3), by performing a univariate rank-based Wilcoxon Rank Sum Test. Using these stringent filters, we identified 1,239 SNPs that were associated with diet-responsive starvation resistance. These SNPs were located within 534 genes (Table S7), 325 of which had human orthologs with a high proportion of the SNPs (>80%) found in non-coding regions (Table S3 and S4). We observed that the largest number of SNPs were associated with HFD (Fig. 2B) and only 7 SNPs affected starvation resistance across all three diets (Fig 2. B). In order to assess the SNP data, we looked at the genomic distribution of all SNPs, including those with predicted diet interactions (Fig. 2C, black dots). To observe the genomic distribution of SNPs on each diet we generated Manhattan plots and observed a higher density of diet-responsive SNPs around chromosome 2L and 3R in flies fed HFD, while HCD and HPD had fewer associated SNPs (Fig 2, D-F black dots). Next, Quantile-Quantile (Q-Q) plots showed an enrichment of significant SNPs compared to the expected p-value distribution, taking into account the genomic population structure of the DGRP, and the correlation structure of diet survival phenotypes (Fig. 2G). We observed no significant associations between the phenotype and covariates (Wolbachia status and large genomic inversions) (data not shown). Thus, we conclude that neither population structure nor additional covariates such as Wolbachia status affected the outcome of our SNP analysis. Overall using conventional SNP analysis resulted in an enrichment of diet-responsive SNPs.

**Figure 2:**
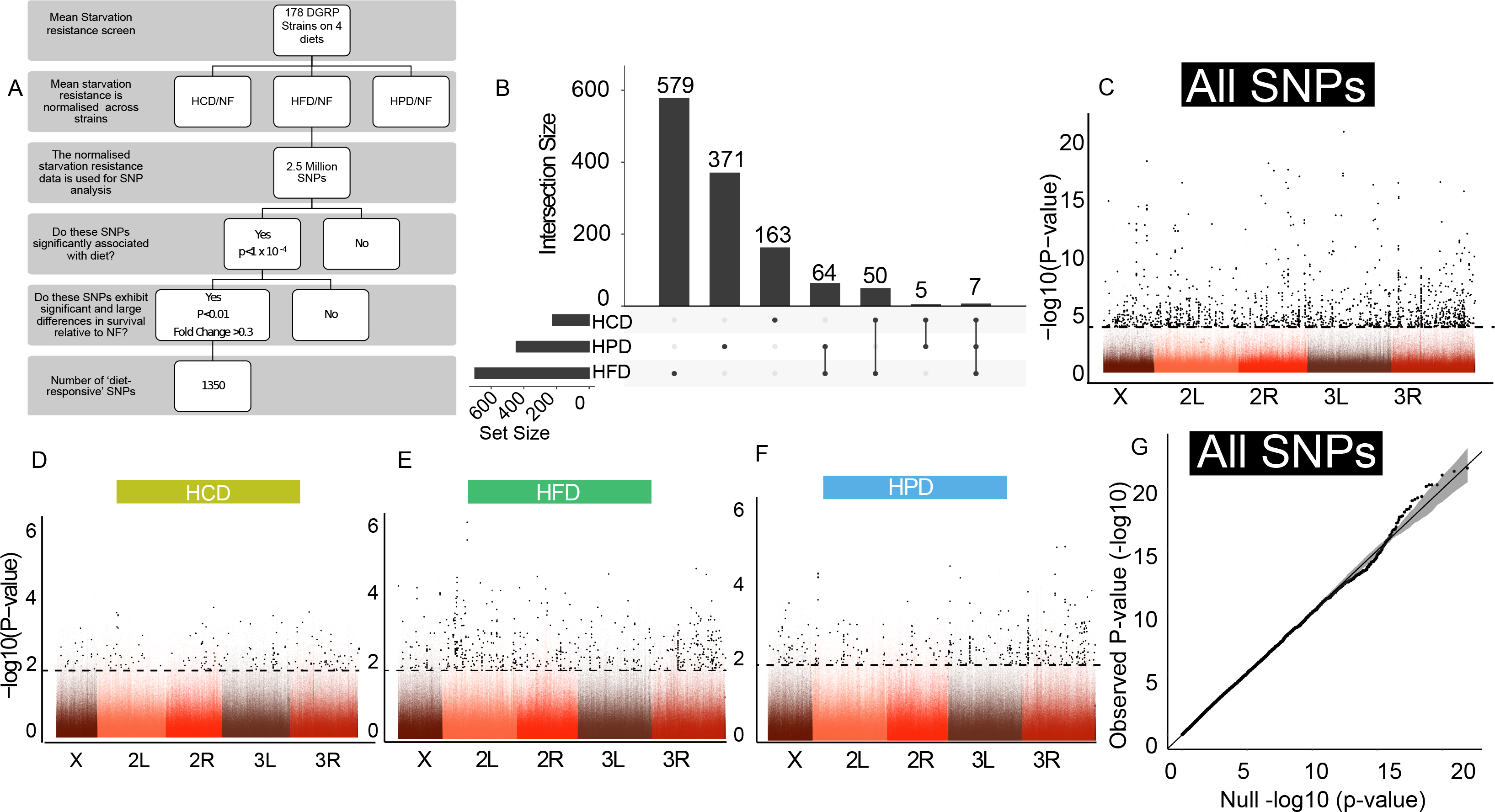
Mapping the SNPs in diet-responsive genes. (A) Schematic of SNP analysis used in this study. (B) An upset plot indicating the degree of overlap of the numbers of highly significant SNPs between each diet and the number of SNPs per diet. (C) Manhattan plot of selected SNPs based on a multi-variate significance p-value of < 1× 10 −4, filtered by a univariate significance of p< 0.01 and a fold change of >0.3. Highly significant SNPs with a univariate p-value <0.01 and a fold change >0.3 are in black. The y-axis in the Manhattan plot is −log (P-value) for the multivariate test. (D-F) Manhattan plots of selected SNPs based on a multi-variate significance p-value of < 11× 10 −4, filtered by a univariate significance of p< 0.01 and a fold change of >0.3 for each diet. Highly significant SNPs with a univariate p-value <0.01 and a fold change >0.3 are in black. G) Quantile-Quantile plots of the distribution of significant MANOVA p-values for all SNPs.

We next undertook a pathway analysis of genes containing significant diet-responsive SNPs using gene ontology (GO) analysis on the human orthologs of the ‘diet-responsive genes’ (Table 2). This revealed stark differences in enriched biochemical processes between the different diets. For instance, genes associated with enhanced starvation resistance (‘up’) on HFD were enriched for peptidases in proteolytic pathways. In contrast, genes associated with reduced starvation resistance (‘down’) after HFD and HPD were enriched for signal transduction and for HFD, the MAPK pathway in particular (Table 2). Meanwhile, extracellular matrix pathways and pathways linked to sulfotransferases were respectively linked to enhanced and reduced starvation resistance after HCD feeding (Table 2). This suggests that individual diet responsiveness is controlled at the pathway level rather than at the level of individual genes *per se*. These data provide an abundant and valuable resource of diet-responsive genes and pathways.

**Table 2:** GO term analysis of human orthologs of diet-responsive genes.

| **HPD "Down"** |  |  |  |  |  |  |
| --- | --- | --- | --- | --- | --- | --- |
| **GO:ID** | **p.value** | **adj.p.**<br>**value** | **Genes In** | **Genes**<br>**Out** | **TERM** | **ONTOL**<br>**OGY** |
| GO:0004888 | 1.84E-08 | 1.07E-04 | 17 | 143 | transmembrane signaling receptor activity | MF |
| GO:0038023 | 1.78E-07 | 3.44E-04 | 19 | 213 | signaling receptor activity | MF |
| GO:0060089 | 1.57E-07 | 3.44E-04 | 20 | 234 | molecular transducer activity | MF |
| GO:0005<br>272 | 9.34E-07 | 1.35E<br>-03 | 6 | 12 | sodium channel<br>activity | MF |
| GO:0015<br>144 | 2.22E-06 | 2.57E<br>-03 | 5 | 7 | carbohydrate<br>transmembrane<br>transporter activity | MF |
| GO:0005<br>887 | 6.56E-06 | 6.33E<br>-03 | 22 | 357 | integral component<br>of plasma<br>membrane | CC |
| **HFD "DOWN"** |  |  |  |  |  |  |
| **GO ID** | **p.value** | **adj.p.**<br>**value** | **Genes In** | **Genes**<br>**Out** | **TERM** | **ONTOL**<br>**OGY** |
| GO:0015<br>294 | 7.51E-07 | 1.09E<br>-03 | 9 | 30 | solute:cation<br>symporter activity | MF |
| GO:0038<br>023 | 3.10E-07 | 1.09E<br>-03 | 22 | 210 | signaling receptor<br>activity | MF |
| GO:0043<br>410 | 7.25E-07 | 1.09E<br>-03 | 19 | 168 | positive regulation<br>of MAPK cascade | BP |
| GO:0060<br>089 | 3.78E-07 | 1.09E<br>-03 | 23 | 231 | molecular<br>transducer activity | MF |
| GO:0019<br>199 | 1.25E-06 | 1.44E<br>-03 | 7 | 15 | transmembrane<br>receptor protein<br>kinase activity | MF |
| GO:0015<br>370 | 1.64E-06 | 1.58E<br>-03 | 8 | 24 | solute:sodium<br>symporter activity | MF |
| GO:0004<br>714 | 2.48E-06 | 2.05E<br>-03 | 6 | 10 | transmembrane<br>receptor protein<br>tyrosine kinase<br>activity | MF |
| GO:0034<br>308 | 3.42E-06 | 2.47E-03 | 8 | 27 | primary alcohol<br>metabolic process | BP |
| GO:0004<br>888 | 6.95E-06 | 4.02E-03 | 16 | 144 | transmembrane<br>signaling receptor<br>activity | MF |
| GO:0043<br>408 | 6.91E-06 | 4.02E-03 | 21 | 237 | regulation of<br>MAPK cascade | BP |
| **HFD "UP"** |  |  |  |  |  |  |
| **GO ID** | **p.value** | **adj.p.**<br>**value** | **Genes In** | **Genes**<br>**Out** | **TERM** | **ONTOL**<br>**OGY** |
| GO:0004<br>180 | 5.29E-09 | 3.06E-05 | 5 | 11 | carboxypeptidase<br>activity | MF |
| GO:0008<br>238 | 3.72E-06 | 1.08E-02 | 5 | 50 | exopeptidase<br>activity | MF |
| **HCD "DOWN"** |  |  |  |  |  |  |
| **GO ID** | **p.value** | **adj.p.**<br>**value** | **Genes In** | **Genes**<br>**Out** | **TERM** | **ONTOL**<br>**OGY** |
| GO:0034<br>035 | 2.43E-07 | 7.02E-04 | 5 | 9 | purine<br>ribonucleoside<br>bisphosphate<br>metabolic process | BP |
| GO:0050<br>427 | 2.43E-07 | 7.02E-04 | 5 | 9 | 3'-<br>phosphoadenosine<br>5'-phosphosulfate<br>metabolic process | BP |
| GO:0008<br>146 | 2.32E-06 | 4.29E-03 | 5 | 16 | sulfotransferase<br>activity | MF |
| GO:0030<br>902 | 2.97E-06 | 4.29E-03 | 8 | 76 | hindbrain<br>development | BP |
| GO:0051<br>923 | 5.89E-06 | 6.81E-03 | 4 | 8 | sulfation | BP |

| HCD "UP" |  |  |  |  |  |  |
| --- | --- | --- | --- | --- | --- | --- |
| GO ID | p.value | adj.p.<br>value | Genes In | Genes<br>Out | TERM | ONTOL<br>OGY |
| GO:0005<br>201 | 1.81E-08 | 1.05E<br>-04 | 5 | 38 | extracellular matrix<br>structural<br>constituent | MF |
| GO:0005<br>581 | 5.98E-08 | 1.73E<br>-04 | 4 | 16 | collagen trimer | CC |
| GO:0005<br>604 | 7.12E-07 | 1.37E<br>-03 | 4 | 32 | basement<br>membrane | CC |
| GO:0062<br>023 | 2.65E-06 | 3.83E<br>-03 | 5 | 110 | collagen-containing<br>extracellular matrix | CC |
| GO:0030<br>198 | 5.63E-06 | 5.18E<br>-03 | 5 | 129 | extracellular matrix<br>organization | BP |
| GO:0031<br>012 | 6.27E-06 | 5.18E<br>-03 | 5 | 132 | extracellular matrix | CC |
| GO:0044<br>420 | 5.95E-06 | 5.18E<br>-03 | 3 | 15 | extracellular matrix<br>component | CC |

Tables show the human orthologs of Drosophila genes associated with a predicted decrease (down) or increase (up) in starvation resistance upon HFD,HPD, or HCD exposure. GO ID with a p-value less than $p < 1E-5$ were selected.

### Candidate gene validation

The following three main criteria were used to select genes for further validation: whether the SNPs passed the more stringent fold change cut-off with respect to diet response (>0.3 log2 fold change); the presence of a human ortholog; and an annotated gene function (Table 3). Using these criteria, we selected 30 genes for further validation using the GAL4-UAS system (Brand and Perrimon 1993). We generated whole-body knockdown flies for each gene in the *W1118* background strain, pre-fed them the four diets and used a more precise (low-throughput) method of monitoring starvation with the *Drosophila* activity monitoring system (DAMs)(Pfeiffenberger *et al.* 2010). Candidate genes were considered validated when knocking down the gene yielded a starvation resistance phenotype (Fig. S2). Strikingly, of the 30 genes tested, whole body depletion of 8 candidate genes led to lethality, while 16 of the remaining 22 candidates displayed a significant diet interaction – 62.5% of these were significant according to a cox hazard multivariate analysis (p<0.05, Table 3, Fig. S2). This validated the use of RNAi knockdown as a useful tool and is consistent with the notion that for many but not all of these genes the corresponding SNP is associated with reduced gene expression or loss of function.

**Table 3:**
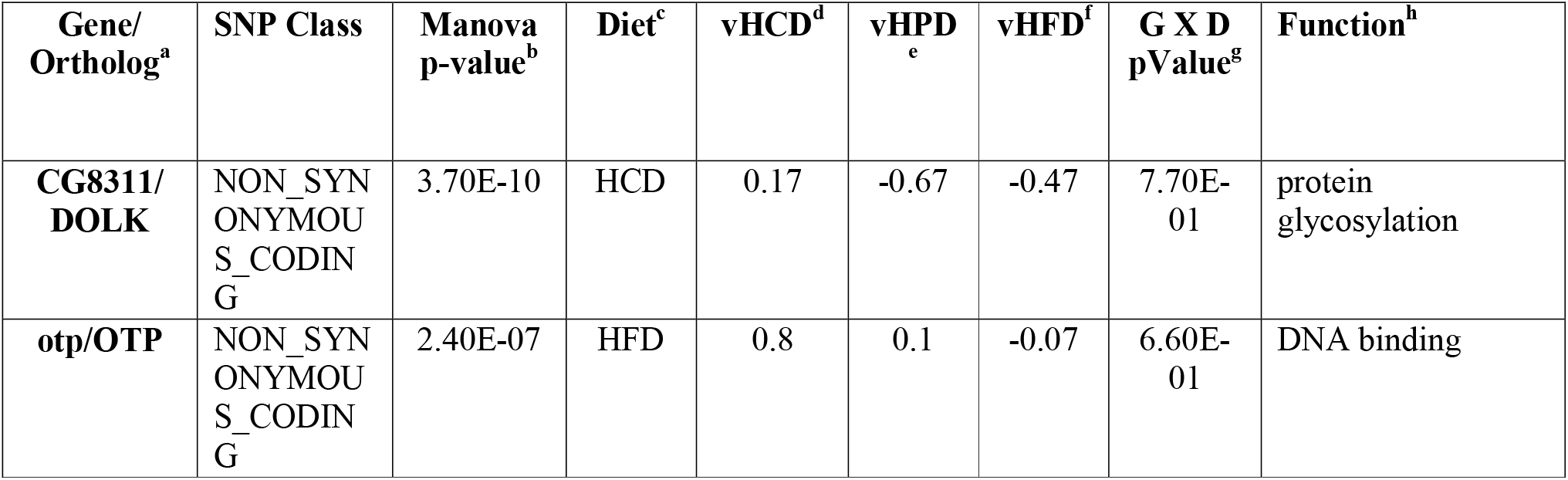
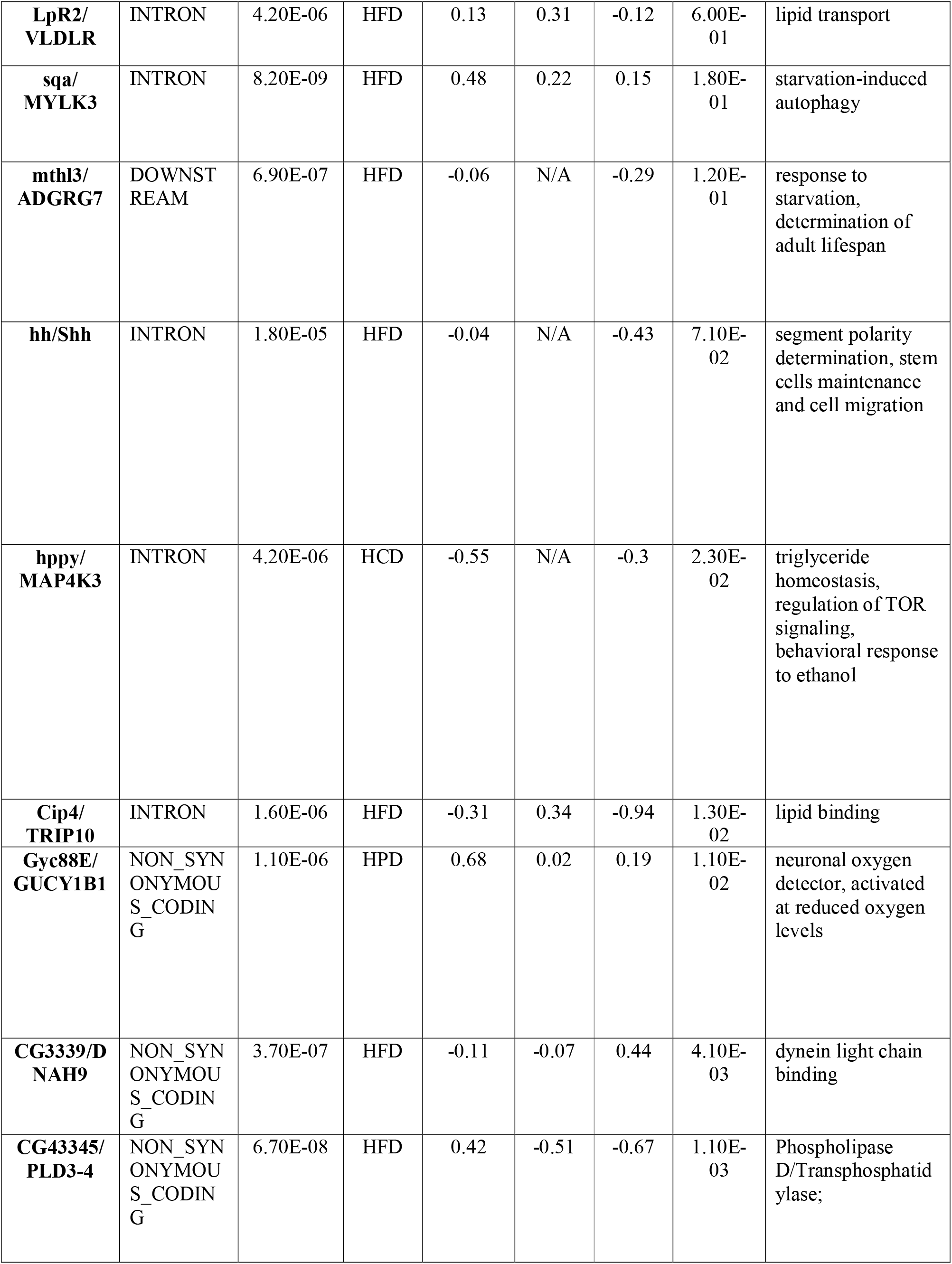
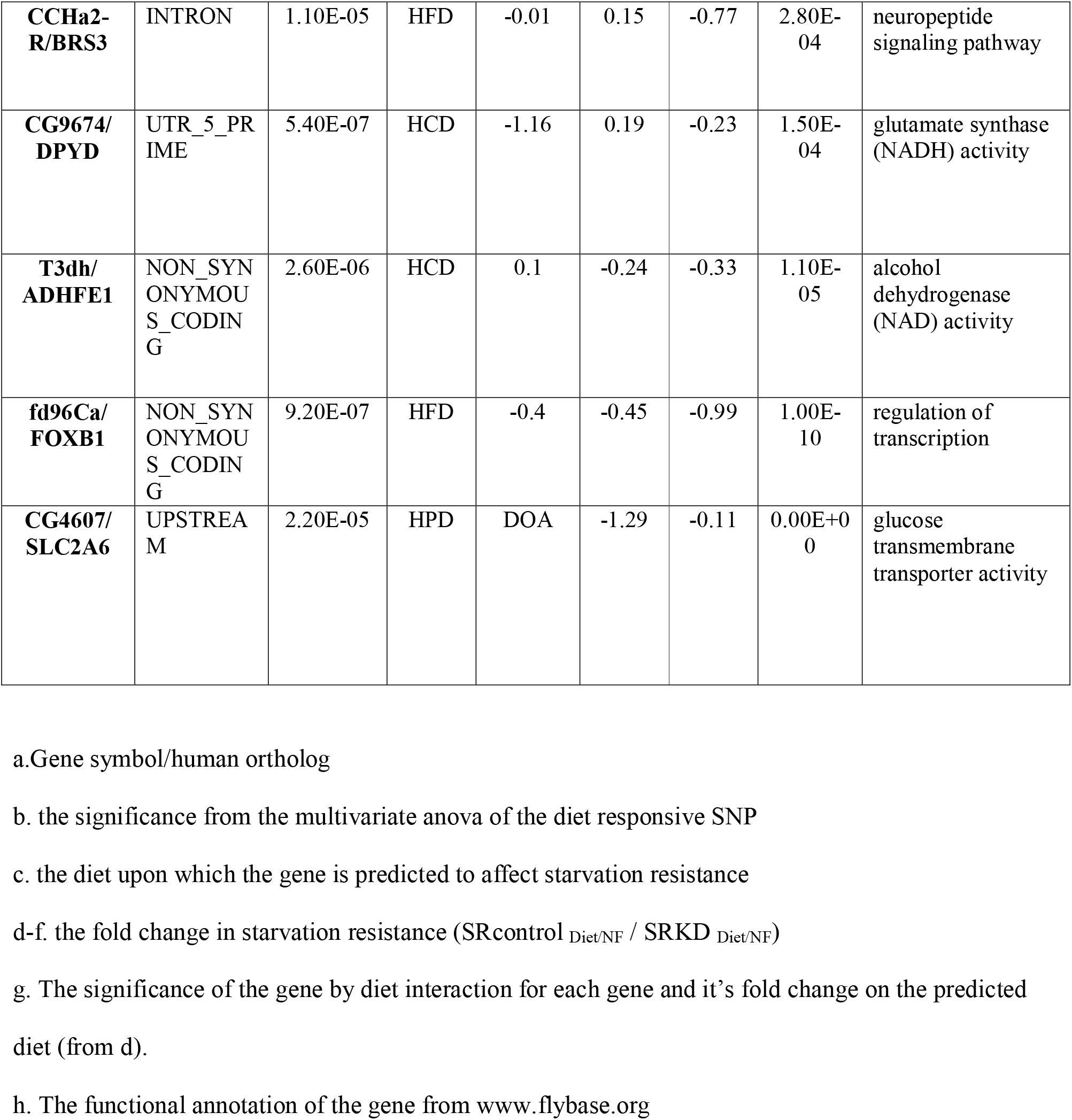
Validated candidate genes.

Of these genes, two were previously associated with starvation resistance: *spaghetti-squash activator* (*sqa*) a myosin light chain kinase that is activated by ATG1 to form autophagosomes during starvation (Tang *et al.* 2011) and *methuselah-like 3* (*mthl3*) known to enhance starvation resistance in the red flour beetle, T. casteneum (Li *et al.* 2014). We also identified a lipid transport protein, *Lipophorin receptor 2 (LpR2)*, required for the transport and binding of lipid molecules from the plasma membrane (Parra-Peralbo and Culi 2011; Rodríguez-Vázquez *et al.* 2015); *Cdc42-interacting protein 4* (*Cip4*), which is implicated in phospholipid binding in *Drosophila* and GLUT4 trafficking in mice (Feng *et al.* 2010; Zobel *et al.* 2015). Furthermore, we identified *CCHamide-2 receptor* (*CCHa-2R*), a G-protein coupled receptor and *Drosophila* ortholog of the bombesin receptor 3 (BRS3), which regulates secretion of *Drosophila* insulin-like peptides in response to nutrient availability (Sano *et al.* 2015). Knockdown of all three genes (*LpR2, Cip4, CCHa-2R*) were associated with increased starvation sensitivity after HFD feeding. Another interesting candidate, *happyhour* (*hppy*), the *Drosophila* ortholog of MAP4K3, a serine/threonine kinase that regulates triglyceride homoeostasis and mTOR signalling (Bryk *et al*, 2010), modulated the response to HCD. Loss of *CG3339*, the *Drosophila* ortholog of dynein heavy chain (DNAH9), a protein that regulates minus end directed microtubule trafficking, enhanced starvation resistance on HCD. The most striking starvation sensitivity in response to diet was observed for *CG9674* (glutamate synthase) and *CG4607* (GLUT6/GLUT8 orthologue). Knockdown of these genes in flies modulated the response to HCD and HPD, respectively (Table 3).

### Analysis of CG4607, an example of a diet responsive gene

*CG4607* was of particular interest because it was starvation resistant on NF, sensitive to HPD, the diet with the lowest amount of sugar, and is homologous to the human glucose transporters SLC2A6 (GLUT6) and SLC2A8(GLUT8). Very little is known about the function of these proteins in mammals and so this raised the possibility that they might play a unique role in the response to sugar. GLUT6 (formerly Glut9) is over expressed in endometrial cancer and is highly expressed in brain, spleen and leukocytes in mice (DOEGE *et al.* 2000). GLUT8 is highly expressed in the testis, heart, brain, liver, fat, and kidneys in mouse and has been shown to respond to insulin and transports trehalose in hepatocytes (Carayannopoulos *et al.* 2000; Mueckler and Thorens 2013; Mayer *et al.* 2016). In flies, *CG4607* is highly expressed in the adult midgut but is modestly expressed in other tissues such as the heart, fat body, salivary gland and fly heads(Robinson *et al.* 2013). However, depletion of *CG4607* with both midgut and fat body drivers did not phenocopy the whole body starvation phenotype (data not shown). Whole body expression of the second hairpin *CG4607 GD3268* was lethal, however, expression of both hairpins using *mef2*-GAL4 was also lethal (data not shown) and suggests that the two hairpins overlap. Remarkably, groups of flies expressing UAS-*CG4607* KK104152 RNAi in the whole body (*CG4607* _KD_) died after only three days of eating HCD (Fig. 3A), while surviving on NF. The DGRP strain 45, which contained the *CG4607* SNP, was also not viable when fed HCD (Table S6), suggesting that the lethality of strain 45 may be due to the presence of the SNP in *CG4607*. To further assess this phenotype, we sought to monitor the behaviour of in single housed control and *CG4607* _KD_ flies while they were eating NF or HCD. We found a striking hyperactivity on HCD that abruptly stopped after 12 h when flies died, as indicated by a total lack of activity (Fig. 3 B, C). The hyperactivity was reminiscent of flies during starvation (Keene *et al.* 2010) and we speculated that the lethality was due to an inability of the flies to consume HCD, thereby experiencing starvation. However, upon measuring caloric intake in control and *CG4607* _KD_ flies we observed that knockdown flies consumed 38% more calories of HCD than controls (Fig. 3D). Thus, the observed hyperactivity of *CG4607*_KD_ flies on HCD may be due to a perception of hunger brought about by the loss of glucose sensitivity, or a defect in the ability to utilise stored nutrients that ultimately lead to lethality.

**Figure 3:**
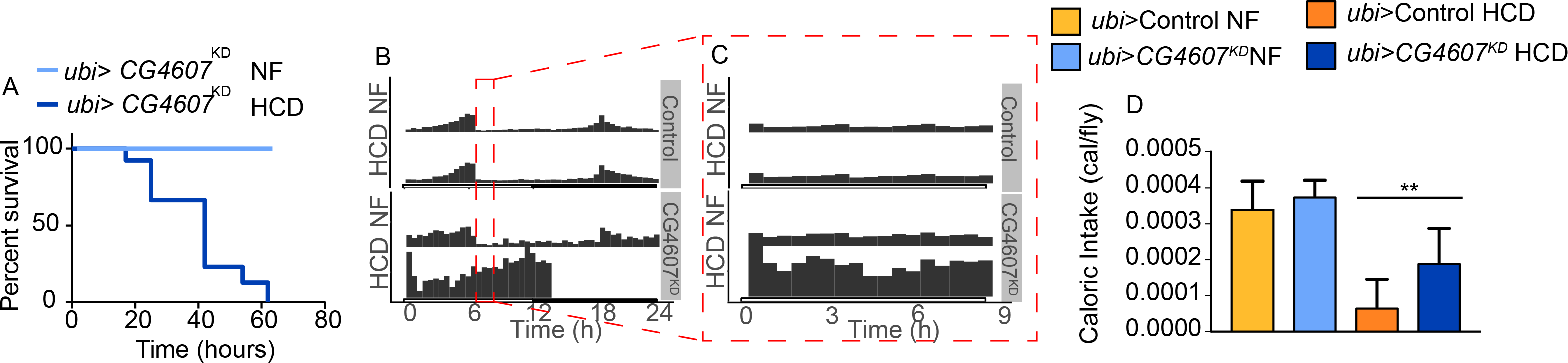
Whole body knockdown of *CG4607* is lethal upon HCD feeding. Survival of *ubi-*GAL4>*CG4607* _KK104152_ flies on NF and HCD. (A) Adult male *ubi-*GAL4>*CG4607* _KK104152_ flies (n=60) were placed on either NF or HCD and manually monitored for lethality every 24 hours. We observed that all *ubi-*GAL4>*CG4607* KK104152 flies were dead within 60 hours after HCD feeding. (B) Actogram of activity and sleep (n=16 flies) wrapped over 24 hours of single housed individual control and *ubi-*GAL4>*CG4607* _KK104152_ flies in DAMS monitors and beam crosses were monitored every 5 minutes while they ate NF or HCD (C) The same actogram as in B, but only the activity over the first 9 hours is shown. Activity data was analysed using the rethomics package in r (Geissmann *et al.* 2019). (D) Caloric intake of adult male flies (n=50/genotype) using the CAFE assay. *ubi-*GAL4>*CG4607* _KK104152_ flies ingested significantly more calories from HCD (p<0.02) than control. Data is mean +/− SD and significance between genotypes was calculated using a t-test, ** p<0.01 (Prism).

Next, we investigated the mechanism by which *CG4607* controlled nutrient storage or utilisation in response to HPD. NF pre-fed *CG4607* _KD_ flies were starvation resistant but became starvation sensitive after becoming low carbohydrate stressed on HPD (Table 3). We posited that the starvation phenotype reflected the levels of energy stores. Fed nutrient levels were similar between *CG4607* _KD_ and control flies on both NF and HPD (Figure 4 A and C), with the exception of TAGs, which were higher in *CG4607* _KD_ animals (Fig. 4 E). However major differences in nutrient storage levels were observed upon starvation on both diets. Fasting of HPD-fed flies resulted in a marked depletion of nutrients in both genotypes with undetectable fasted nutrient levels in control flies. Yet, fasted levels of glycogen, glucose and TAGs were still higher in *CG4607* _KD_ flies than in control flies (Fig. 4 B, D, G). Overall, our data shows that relative to control flies, *CG4607* _KD_ flies have increased caloric intake and excessive energy stores on NF, which are reduced upon HPD feeding. This suggests that although *CG4607* _KD_ flies have an excess of energy stores, these are still mobilised during starvation. However, given that HPD *CG4607* _KD_ flies still die faster than HPD control flies during the starvation resistance assay, this suggests an inefficient mobilisation of energy stores.

**Figure 4:**
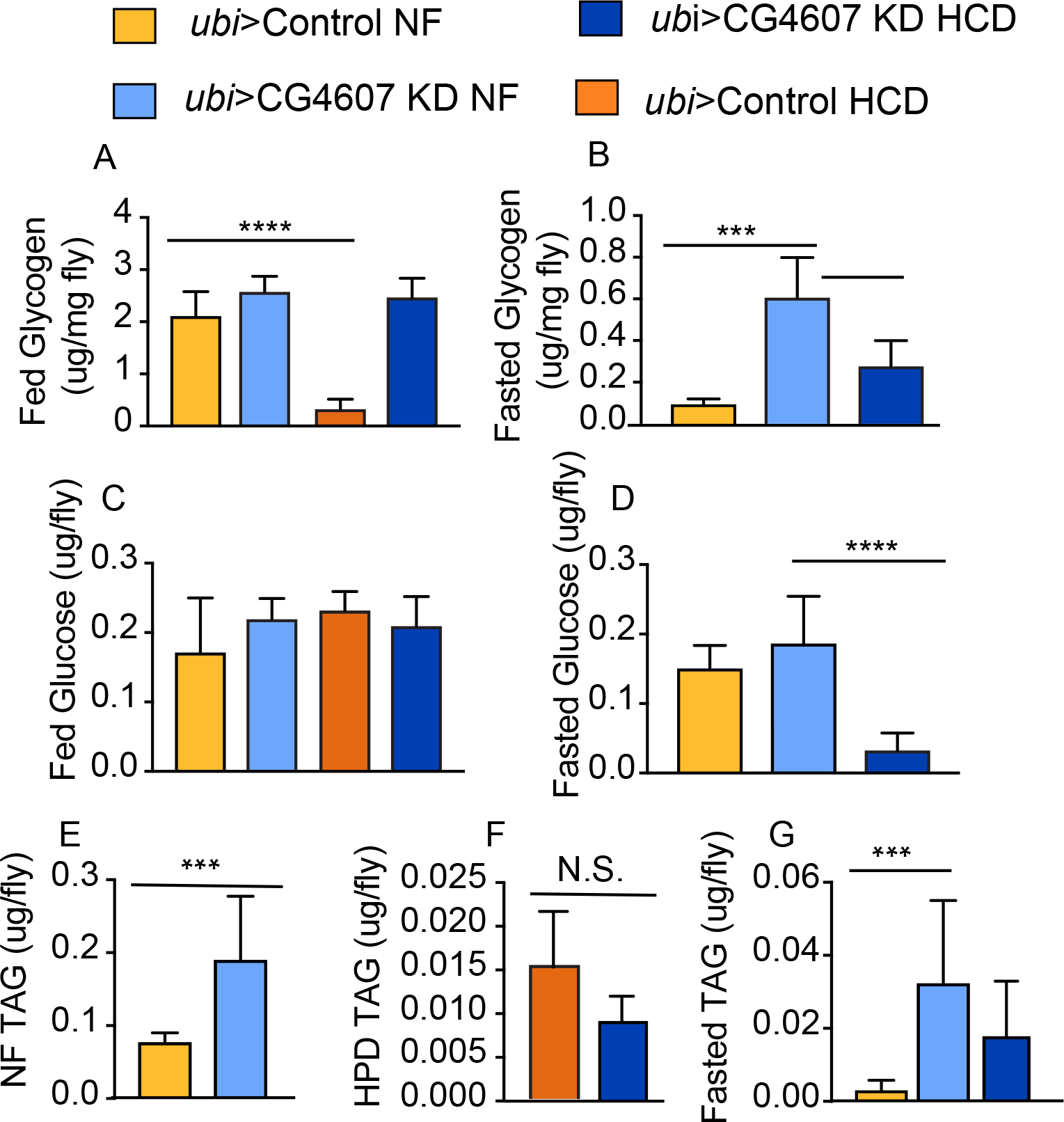
*CG4607* interacts with HPD to regulate starvation resistance. (A) Glycogen content was significantly reduced between control flies fed NF and HPD. However, glycogen content was the same between NF and HPD fed *ubi-*GAL4>*CG4607* _KK104152_ flies (B) Fasted glycogen was reduced in NF fed control flies compared to NF fed *ubi-* GAL4>*CG4607* _KK104152_ flies, but pre-feeding with HPD reduced glycogen levels of *ubi-*GAL4>*CG4607* _KK104152_ flies, while starvation after control flies ate HPD was lethal. (C) Glucose content was not significantly different across genotypes. (D) Fasted glucose levels of NF pre-fed flies had similar glucose levels, however, glucose levels were significantly reduced in HPD pre-fed *ubi-*GAL4>*CG4607* _KK104152_. (E)Triglyceride levels of *ubi-*GAL4>*CG4607* _KK104152_ animals were increased on NF feeding, but reduced upon eating HPD. (F) Fasted TAG stores were reduced in control flies pre-fed NF, while NF fed *CG4607_KD_* flies maintained their TAG stores. (G) HPD pre-fed flies had reduced TAG stores. (all experiments n=36 flies/genotype and representative of two experiments) Significance between genotypes was calculated using a one-way ANOVA, ** p<0.01, *** p<0.001 (Prism).

To further probe the role that *CG4607* plays in mobilising energy stores during starvation, we wanted to explore if CG4607 functions like a glucose transporter. To address this, we measured glucose utilisation by feeding flies an NF diet containing _14_C-radiolabelled glucose and monitoring glucose incorporation into CO_2_ and lipids (Francis *et al.* 2019; Krycer *et al.* 2019). We observed that *CG4607* _KD_ flies exhibited lower levels of glucose oxidation and incorporation into lipids compared to controls (Fig. 5 A, B), indicating reduced glucose utilisation. This was not due to lower food intake in the *CG4607* _KD_ flies (Fig. 5C). Given the effect of depleting *CG4607* on glucose utilisation, we wanted to see where it was localised and posited that its localisation would be similar to its mammalian homologs. The mammalian GLUT6 and GLUT8 both localise to lysosomes (Diril *et al.* 2009; Maedera *et al.* 2019). To determine if the localisation of CG4607 was conserved, we created and expressed CG4607-mRUBY3 in HeLa cells. We used immunofluorescence microscopy to observe that, like GLUT6 and GLUT8, CG4607 partially co-localised (Avg. Pearson’s r = 0.57, n=13 cells) with lysosomal markers (Fig.5 D, E). Taken together, our data show that CG4607, a GLUT6/8 orthologue, is an example of a diet-responsive gene and that responds to dietary sugar by regulating lysosomal glucose metabolism.

**Figure 5:**
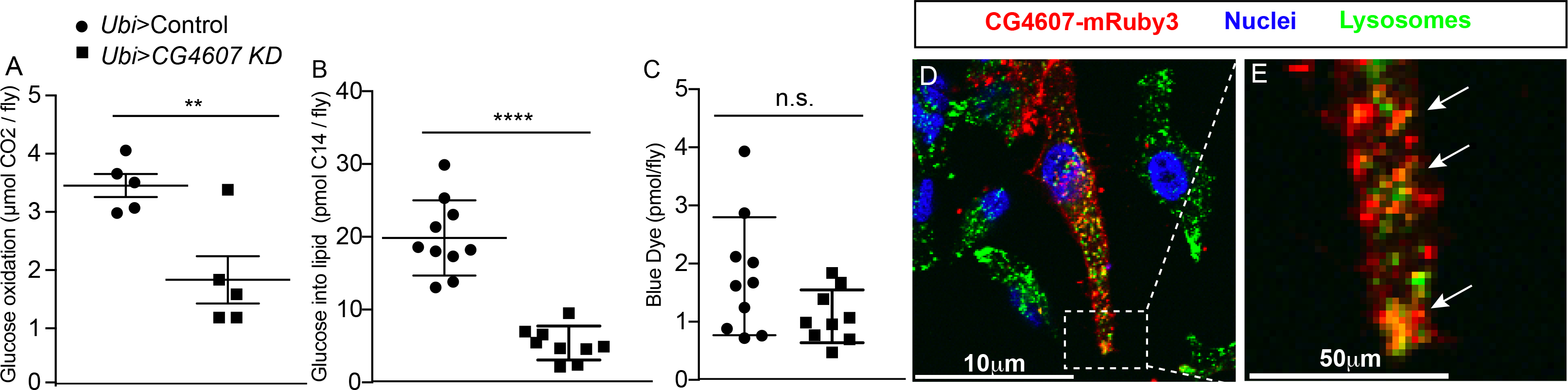
*CG4607* controls glucose utilisation and uptake. (A-C) All the following experiments were performed on n= 40 control and *ubi-*GAL4>*CG4607* _KK104152_ males as previously described (Francis *et al.* 2019). (A) Glucose oxidation is reduced after 4 hours of administrating C14 labelled glucose. (B) C14 incorporation into TAGs is reduced in *CG4607*_KD_ flies. (C) The accumulation of blue dye, which indicates food intake, is not significantly different between control and *CG4607*_KD_ flies during the assay. The data is from two independent experiments. Significance between genotypes was calculated using a student’s t-test, ** p<0.01, **** p<0.0001 (Prism). (D,E) Micrographs of fixed and immuno-stained HeLa cells were transfected with pCMV-CG4607-mRuby3 (red), immunostained with anti-LAMP1 antibody (green) and counterstained with Hoechst (blue). (D, inset (E)) CG4607-mRuby3 colocalises with Lamp1 expressing vesicles.

## Discussion

The DGRP is a powerful tool for understanding the genetics driving variation of metabolic phenotypes (Mackay *et al.* 2012; Garlapow *et al.* 2015; Unckless *et al.* 2015; Nelson *et al.* 2016). Our approach measured starvation resistance in the adult male population after exposure to diets that vary in sugar, protein and fat content. Using this method, the DGRP supported the notion that nutrition responses vary considerably between individuals of different genetic background. Although, diet had a major influence on starvation resistance, the diets that enhanced or reduced starvation resistance differed markedly between *Drosophila* strains. In particular, exposure to diets high in sugar (HCD) or protein (HPD) elicited a greater genetic contribution to phenotypic variation. These data highlight a heterogeneity in the response to diet that underlies the fundamental principles of personalised nutrition. The SNPs we identified provide a resource dataset for further study. Out of 39 candidate diet responsive genes that were selected for more detailed analysis, we validated 21 genes as bona fide diet responsive genes. Knock down of one such gene, *CG4607*, was linked to increased caloric intake and nutrient storage combined with reduced glucose utilisation.

In relation to our study, since the natural food sources of *Drosophila* are rotting fruits that contain sugar and yeast (Markow 2015), we postulate that genetic variants involved in the processing, storage and utilization of sugar or protein (from yeast) contribute to differences in starvation resistance. Indeed, changes in sugar and lipid handling enzymes have previously been linked to starvation resistance (Harshman *et al.* 1999). Alternatively, sugar and protein (from yeast) are dominant drivers of food intake with major sensory systems to control intake of these macronutrients and hence these are more likely to be under genetic control (Chng *et al.* 2017; May *et al.* 2019). Overall, this demonstrates that the mechanisms that underpin diet sensitivity are complex and involve differences in food preference and thus food intake as well as differences in nutrient metabolism (energy storage and utilisation), all of which are regulated at least in part by one’s genotype.

This study provides a rich resource of diet responsive genes and pathways. The majority (>80%) of the SNPs in candidate genes were non-coding, a finding that is consistent with previous DGRP (Mackay *et al.* 2012) and human GWAS (Gallagher and Chen-Plotkin 2018) and the functional consequences of such SNPs remain to be identified. As a proof of principle, we were able to validate candidate genes with intronic SNPs using RNAi knockdown indicating that these types of mutations may control expression of these genes. However, further studies examining transcriptional control are required to validate this conclusion.

We focused on *CG4607*, a validated diet-responsive gene. The SNP in *CG4607* was non-coding and within the 5’-UTR, suggesting that it could affect splicing or post-transcriptional processing. Additionally, the SNP contains a regulatory transcription factor binding site for the *invective* transcriptional repressor. While it may be of interest to determine if the SNP alters the expression of CG4607, our validation using RNAi knock down clearly shows that CG4607 is a diet-responsive gene. SNPs in *CG4607* were identified in a GWAS of sleep/activity (Harbison *et al.* 2013). Our study provides several lines of evidence that CG4607 mediates its effects via regulating glucose metabolism. First, whole body depletion of *CG4607* resulted in lethality after 3 days on a high sugar diet (HCD) (Fig. 3A) and the lethality was accompanied by hyperactivity and an increase in caloric intake on HCD compared to control flies. This is symptomatic of a starvation phenotype (Yu *et al*, 2016; Yang *et al*, 2015). Second, *CG4607 _KD_* animals have higher energy stores on NF compared to control flies, likely due to impaired glucose utilisation. Third, our metabolic labelling experiments show that *CG4607 _KD_* animals exhibit reduced glucose utilisation.

Last, the closest mammalian orthologues of *CG4607* are the human glucose transporters GLUT6 and GLUT8. Recently, GLUT8 has been shown to transport trehalose in mammalian cells (Mayer *et al.* 2016) suggesting that CG4607 may play a similar role in flies. However, loss of *CG4607* does not phenocopy elevated or reduced trehalose levels and may not play a major role in trehalose transport(Matsuda *et al.* 2015; Yasugi *et al.* 2017). Furthermore, GLUT8 regulates AMPK phosphorylation and signalling via trehalose transport (Mayer *et al.* 2016; Narita *et al.* 2019) and AMPK deficient flies are starvation sensitive (Johnson *et al.* 2010). It would be of interest to determine if CG4607 regulates AMPK in a similar manner as its mammalian ortholog. Given that AMPK is an energy sensor perhaps energy sensing pathways play a part in diet-dependent differences in starvation resistance.

Like these GLUT transporters, CG4607 is also targeted to lysosomes instead of the plasma membrane, unlike other facilitative sugar transporters (Lisinski *et al.*; Maedera *et al.* 2019). This is intriguing as this co-localises with mTORC1 (Lee *et al.* 2009; Efeyan and Sabatini 2013), which regulates numerous metabolic processes including glycogen breakdown in autophagic vesicles (Mony *et al.* 2016; Zhao *et al.* 2018). Given that compared to control flies, *CG4607 _KD_* flies exhibited higher glycogen levels after starvation, and broke down more TAGs during starvation, the metabolic phenotype of *CG4607 _KD_* flies resembles a lysosomal glycogen storage disease, where lysosomal glycogen cannot be accessed for energy utilisation in the cytosol. Together, this provides strong evidence that CG4607 mediates its effects on starvation resistance by regulating glucose metabolism. Hence this highlights the importance of glucose metabolism as a potential diet responsive pathway.

Overall, our findings demonstrate that gene-diet interactions are an impact factor to consider in metabolic homeostasis. This has substantial ramifications for human health because it means that the concept of a ‘healthy’ diet varies between individuals, thus questioning population-wide nutritional recommendations. While our study provides the basis for a nutrigenomics initiative such an endeavour is likely to require a substantial future investment at the clinical level.

## Materials and Methods

### Drosophila Stocks and procedures

#### Stocks

DGRP (Bloomington *Drosophila* Stock Center, Indiana, USA), RNAi reagents (CG4607 v107219 and v5450, VDRC, Vienna, Austria), *ubiquitous*-GAL4 (Bloomington # 32551) and *CG*-Gal4 (Bloomington # 7011). Flies were maintained at standard temperature and 12h light/dark cycle. DGRP flies were expanded in bottles before collecting adult males for experiments. 5 replicates of ten 3-5 day old adult males from each strain were collected and passaged onto each diet. Food was changed every other day, and the mortality rate was monitored for the ten days of diet treatment. The diets were well tolerated with a similar lethality on HFD during pre-feeding compared to other diets and did not affect the starvation assay. Afterwards, males were placed into starvation vials with kimwipes and 1 mL of water and monitored every 12 hours for death. We initially tested 5 diets (NF, HCD, HFD, HPD and CR). However, we discovered that the CR diet (6.5% sugar and yeast) was not very different compared to NF. Thus, we chose to focus only on NF, HCD, HPD and HFD. The median and mean starvation resistance was analysed using Prism and the R (CRAN, survival and survminer packages).

### Heritability estimates using LMMs

We estimated heritability from two LMMs of log survival. Model 1 included diet as a fixed effect and a random intercept for genotype. Model 2 also included a random slope for diet effects among genotype. Heritabilities were calculated as σ_2Genotype_ / (σ_2Genotype_ + σ_2Residual_) × 100. For model 2, diet specific σ_2Genotype_ was calculated as σ_2Intercept_ + σ_2Diet_ + 2 × ρ_Intercept_ × σ_intercept_ × σ_Diet_. A likelihood ratio test suggested model 2 gives significantly better fit to the data than model 1.

### Statistical SNP analysis

In order to determine the diet-responsive SNPs we performed bioinformatic analysis on mean starvation resistance data of each DGRP strain. To use the effect size of the diets, the mean starvation resistance of DGRP strains on HCD, HFD and HPD was divided by the mean starvation resistance on NF. This normalised starvation resistance was fed into the analysis pipeline. DGRP SNP genotypes were downloaded from the DGRP Freeze 2 online resource http://dgrp2.gnets.ncsu.edu/data.html for all lines considered in this manuscript. SNPs were filtered so at least five lines contained one of each of the reference and alternate alleles, resulting in testing across 2,455,135 SNPs. Lines with missing allele information for a given variant were not considered within each test. Statistical testing included multivariate analysis of variance (MANOVA) testing, with Wolbachia status as a covariate per variant, with an unadjusted P-value < 1e-4 as significant, as well as Wilcoxon Rank Sum Tests per diet. The overall enrichment of the multivariate response was assessed by comparison to the null p-value distribution under random permutation of line labels, thereby retaining the overall phenotype and SNP-SNP correlation structure (Fig 2G). To assess the effect size, we calculated the median difference in phenotype between ‘reference’ and ‘alternate’ allele groups per diet, phenotype being the log2 ratio of survival to NF per line. We selected diet-SNP pairs for further consideration if they were significant for both the MANOVA test and Wilcoxon Rank Sum Test, and had an absolute difference in median phenotypes of at least 0.3 (log2 fold change).

### Gene ontology analysis

Gene ontology and pathway analyses were performed using Fisher’s Exact Test in the ‘goseq’ R package. We used human orthologs for the analysis since GO analysis of fly genes led to very broad non-specific categories (results not shown). Furthermore, using only the human orthologs would limit us to conserved genes. A total of 6,056 pathways, based on human genes, with at least 10 and at most 500 genes were kept, with a ‘gene universe’ of 7,677 human genes that are homologues of the fly genes tested. Pathways were considered significant if FDR-corrected values were below 0.05, per diet and direction of phenotype change.

### Validation/Automated starvation resistance (DAMS assay)

We used the GAL4-UAS gene expression system to validate the 30-candidate diet-responsive genes. RNAi knockdown fly lines (see table S5 for reagent ID) were mated to 20 *ubi*-Gal4 females. Sixteen 3-5 day old males were placed on five different diets for ten days. The food was changed every other day for ten days until males were placed into the DAMS apparatus (Trikinetics, inc., USA). The flies were loaded into DAMS tubes containing 2% agar and monitored every 5 minutes for starvation resistance. Candidate genes were considered validated when the fold change in survival on each diet (relative to NF) corroborated with the SNP analysis (Table 3, Table S5). The gene by diet p-values were generated using the Cox proportional hazards regression model (survival package, R).

### Measuring Activity

To measure fly activity while eating NF or HCD we placed 3-5 day old male flies into DAMs tubes with either NF or HCD and monitored for activity for ten days. Since *CG4607 _KD_* flies died within the first 24 hours after being placed on HCD, we analysed the activity data from the beginning of the monitoring. DAMS data were analysed using the R survminer and rethomics (Geissmann *et al.* 2019) package.

### Blue Dye Extraction and measurements

To determine the amount of food ingested during the gas-trap assay, we extracted and measured blue dye that was mixed with the radiolabelled food. In short term experiments, the blue dye serves as a proxy for food intake (Krycer *et al.* 2019). 4 replicates of ten flies were collected and homogenized (Reche MM400) in 100uL of water. Samples were briefly spun down and the supernatant was dried down in the Genevac personal evaporator. Dried samples were reconstituted in 50uL of water, vortexed and placed into a 96 well plate for measurement at 628nm in a spectrophotometer. Dilutions of blue dye (Queenie Brand, Coles Supermarket, Australia) were used as a reference. Data was analysed in Excel (Microsoft) and plotted in Prism (Graphpad). Statistical significance was calculated between genotypes using Student’s t-test.

### Capillary feeding assay

We used the Café assay as previously described (Lee *et al.* 2008) to determine the amount of the experimental diet eaten. 10 replicates of five 3-5 old adult males were placed into vials with water soaked kimwipes and sealed with a rubber stopper with two holes. 5uL capillary tubes with food was placed through one hole and into the vials to allow flies to feed for 24 hours(Lee *et al.* 2008). The diets were composed of yeast (MP Biomedicals cat # 2232731), and sucrose (Table Sugar, Coles Supermarket, Australia).

### Experimental Diets

The experimental diets that were used throughout this study were made up of agar (Sigma) and torula yeast (H.J Langdon & Co, Victoria, Australia), Sucrose (Table sugar, Coles, Victoria, Australia) and extra virgin coconut oil (Absolute Organics, NSW, Australia).

### Gas Trap Assay to measure CO2 and Triglycerides

We used a Gas Trap Assay to assess the capability of *CG4607 _KD_* flies to utilise glucose. The gas trap protocol has been previously described(Francis *et al.* 2019). Briefly, 4 replicates of ten male adult 3-5-day old flies were starved overnight with a Kim wipe and 1mL of water. Flies were placed into 12 well plates containing glucose radiolabelled food and blue dye. We measured glucose oxidation and processed the flies for TAGs and blue dye content as described.

### Triglycerides Assay

To measure triglyceride content we used a triglyceride extraction method performed as previously described (FOLCH *et al.* 1957). 6 to 10 replicates of six 3-5 day old flies were collected and washed in 4 dilutions of isopropanol to remove excess food. The lipids were collected after extraction, evaporated under N2 gas and reconstituted with 95% ethanol. Scintillant was added to samples with radioactive tracer instead of ethanol and read on a beta counter (Beckman Coulter). For non-radioactive samples: samples were spun and placed into 96 well plates (Sigma-Aldrich, # CLS9018BC) and incubated with triglyceride reagent (200uL; Thermo Fischer Cat #TR22421) for at 37C for 30min. Precimat glycerol reagent (Thermo Fischer # NC0091901) was used as a reference. Total absorbance at 500 nm was measured in a plate reader (Beckman) and subtracted from a blank before determining the amount of triglyceride using the reference standard curve. All calculations were performed in Excel (Microsoft) and graphed in Prism.

### Glycogen Assay

To determine the amount of glycogen we collected 6 replicates of six male flies and washed them in several dilutions of isopropanol to remove food. Fasted flies were collected after 24 hours of starvation. Flies were homogenised in 1M KOH for 30 seconds using steel balls and a tissue lyser (Resche MM400). Samples were heated for 30 min at 70 degrees C. Saturated Na2SO4 was added following 95% Ethanol for precipitation. The pellet was spun down and then reconstituted in water, heated at 70 degrees and 95% Ethanol was added again. The pellet was spun down, and amyloglucosidase (Merck # A7420) were added overnight at 37 degrees. Samples were spun and placed into 96 well plates (Sigma-Aldrich, # CLS9018BC) and incubated with glucose oxidase reagent (200uL; Thermo Fischer, TR15221) for at 37C for 30min. 1mg/ml glucose was used as a reference. Total absorbance at 500nm was measured in a plate reader (Beckman) and subtracted from a blank before determining the amount of glucose using the reference standard curve. All calculations were performed in Excel (Microsoft) and graphed in Prism.

### Glucose assay

Glucose was measured from the aqueous phase of the triglyceride extraction (see above): the aqueous mixture was evaporated in a Genevac E2-3 evaporator until a dried pellet was visible. The pellet was reconstituted with water and glucose was measured as described for glycogen.

### Generation of CG4607-mRuby3

In order to determine whether the localisation of CG4607 was conserved we created a plasmid to express CG4607 in mammalian cells. CG4607-mRuby3 construct was created through Gibson cloning (Gibson *et al.* 2009). The CG4607 cDNA (clone RH58543 #`11058`, Drosophila Genome Resource Center, Indiana, USA) was PCR amplified using the following primers:

dCG4607GibF1: GGACTCAGATCTCGAGACAAGATGAAGGGCCAGCAGGAGGAG
dCG4607GibR1: CATGCTGCCttCAGCTGAGGACAATTTCTTTAGGAACACTT
The backbone GLUT4-mRuby3 was PCR amplified to include overhangs using the following primers: mRuby3GibF1: TCCTCAGCTGAAGGCAGCATG
mRuby3GibR1: AGCTGAGGATCCCTTGTCTCGAGATCTGAGTCC
PCR products were placed together with Gibson master mix, and the resulting plasmid was sequenced before cell transfection.

### Cell Culture and Immunostaining

In order to observe the localisation of CG4607 we expressed an ectopic construct, encoding CG4607 fused to mRuby3, in mammalian HeLa cells. HeLa cells were kept in DMEM with 1% glutamax and 10% FCS at 37 degrees and 5% CO2. Cells were transfected with lipofectamine 2000 and split onto coverslips at two ×10^5^ cells/mL. To determine the localisation of CG4607 with lysosomes we used immunostaining with anti-LAMP1 antibodies and then looked at colocalization between the lysosomal marker and CG4607-mRuby3. Coverslips were fixed in 4% paraformaldehyde, washed with PBS, and blocked for 30 minutes with 0.02% saponin (Sigma,) and 2% BSA in PBS. Primary antibodies: ms anti-LAMP1(1:100, Developmental Studies Hybridoma Bank 4C CR). Secondary antibodies: (Gt anti-mouse 488 (1:200, Invitrogen)). Coverslips were mounted in mowiol and imaged using a 60X water objective on the A1R confocal (Nikon). Colocalisation analysis was performed using the coloc2 plugin in Fiji (Schindelin *et al.* 2012) ImageJ, NIH, Bethesda, MD).

### qPCR of knockdown

We wanted to determine the degree of whole body depletion of CG4607. We used qPCR to look at the number of transcripts in control and CG4607 _KD_ animals. Three replicates of 10 flies were homogenized in TRIzol™ Reagent (Invitrogen, 15596026) and RNA was precipitated out. cDNA was created (superscript II, Invitrogen 18064014). Tubulin was used as a housekeeping gene and the following primers were used to amplify CG4607:

CG4607 F3 ACTCCCACGCGAAGGAGAA
CG4607 R3 GCTGATTGAGAGTAACTGCCG

The samples were run using the ROCHE Lightcycler 480 II (Roche). The knockdown efficiency was calculated using the delta-delta Ct method (Excel and Graphpad, Prism) and the Ct values were graphed. Significance between the control and knockdown transcript was calculated using a student’s t-test (p<0.0001 ****).

## Acknowledgements

The authors would like to thank Elise Needham for edits, Roel Bevers and Bart Deplancke for useful suggestions and members of the James Lab for helpful discussions. The authors acknowledge the facilities, and the scientific and technical assistance, of the Australian Microscopy & Microanalysis Research Facility at the Charles Perkins Centre, The University of Sydney.

